# Organism-wide spatiotemporal profiling of gene expression utilizing a X-CreERT2/Ai9 tracing system

**DOI:** 10.1101/2025.07.03.662901

**Authors:** Feiyun Li, Qing Yao, Mingjue Chen, Shuangshuang He, Yi Ran, Jianglong Li, Lijun Lin, Guozhi Xiao

**Author notes:** Correspondence (G. X.).

## Abstract

This study utilizes a novel inducible X-CreERT2/Ai9 system to comprehensively delineate the spatial expression landscape of *Fermt2*, which encodes a key focal adhesion protein Kindlin-2, across organs in mice. We can confirm all known Kindlin-2-expressing cells and identify any previously unknown Kindlin-2-expressing cells and those that do not express Kindlin-2 in all tissues/organs in mice. We have developed this cost-effective, rapid, and high-fidelity system and implemented its first whole-organism application for comprehensive gene expression profiling. This work not only provides unprecedented insights into roles of Kindlin-2 in diverse physiologies and pathologies, but also establishes the inducible X-CreERT2/Ai9 system as a powerful paradigm-shifting platform for organism-wide pan-organ spatiotemporal expression profiling of any target genes encoding any proteins and micro-RNAs. By enabling high-resolution mapping of drug target expression, this system may lay critical groundwork for elucidating previously unknown adverse and beneficial effects of targeted therapies, offering transformative potential for precision medicine.

## INTRODUCTION

Contemporary biomedical research increasingly prioritizes targeted therapies^1,2^. Currently, comprehensive atlases of target gene expression at the pan-organ level are still lacking. This obscures therapeutic windows and unexplored indications for both existing and emerging targeted therapies. Therefore, establishing a new methodology for systematic delineation of target gene expression patterns under physiological and pathological conditions is imperative for optimizing current regimens and developing next-generation precision therapeutics.

Current knowledge regarding expression of any target genes is basically from individual studies by utilizing conventional technologies, such as in situ hybridization (ISH), immunofluorescence (IF), and emerging spatial transcriptomics. While results from using these technologies provide valuable spatial gene expression information, they have major limitations. ISH/IF require high-affinity, high-specificity probes/antibodies whose development, validation, and implementation are highly time-consuming and technically impossible to perform at the pan-organ level. Furthermore, cross-reactivity or epitope masking may yield false-positive and false-negative results, compromising analytical reliability and reproducibility. While single-cell sequencing technology can identify cell populations expressing target genes, it cannot provide their spatial expression information. It is true that spatial transcriptomics achieves transcriptome-wide spatial resolution within biological specimens; its prohibitive cost essentially impedes its large-scale and high-throughput applications, especially at the pan-organ/tissue level.

The inducible CreERT2/Ai9 system is a new technology that has been mainly used for lineage tracing studies^3–6^. Its exceptionally high spatiotemporal controllability and specificity make it a possible versatile genetic toolkit for systemic expression mapping. In this study, we have developed a novel transgenic mouse model by inserting a 2A-CreERT2 cassette at the 3’-end of exon 15 in the *Fermt2* gene, which encodes Kindlin-2, thus coupling spatiotemporal expression of Kindlin-2 with that of CreERT2. Tamoxifen (TAM)-induced recombination triggers tdTomato expression exclusively in Kindlin-2-expressing cells, which allows rapid and systematic detection of Kindlin-2 expression patterns in multiple tissues/organs. Following TAM induction in *Fermt2-CreERT2/Ai9* mice, we can rapidly visualize spatial distributions of Kindlin-2-expressing cell populations with high precision. This approach delivers major advantages, including speed, high-fidelity and high cost-effectiveness, which can be performed at the pan-organ level. Importantly, reporter expression is precisely governed by the endogenous promoters of target genes, thus eliminating specificity concerns inherent to other technologies. This approach establishes a novel paradigm for systemic gene expression profiling of *Fermt2* and any other genes encoding proteins and miRNAs and provides a conceptual framework to guide subsequent investigations into their functions under physiological and pathological conditions across diverse organ systems.

## RESULTS

### Construction of the inducible *Fermt2-CreERT2/Ai9* genetic tracing system

To establish an inducible genetic tracing system for Kindlin-2-expressing cells, we employed CRISPR/Cas9-mediated homologous recombination to precisely insert a 2A-CreERT2 cassette at the 3’ terminus of exon 15 in the *Fermt2* locus, generating inducible *Fermt2-CreERT2* knock-in mice (Supplementary Figure 1A). Subsequent crossbreeding with *Ai9* reporter mice yielded double-allele *Fermt2-CreERT2 Ai9^fl/+^* progeny. In this model, TAM (100 mg/kg, i.p.) administration induces nuclear translocation of CreERT2, mediating excision of the loxP-Stop-loxP cassette in *Ai9* reporter cells. This recombination event triggers cell-autonomous tdTomato expression, yielding intense orange-red fluorescence specifically in Kindlin-2-positive cell populations (Supplementary Figure 1B). Importantly, neither genetic modifications nor TAM administration caused marked abnormalities in appearance (Supplementary Figure 1D).

For systemic mapping of Kindlin-2 expression patterns, 2-month-old *Fermt2-CreERT2 Ai9^fl/+^* mice underwent TAM induction followed by 30-day lineage tracing (Supplementary Figure 1C). Various organs were collected and processed with standardized tissue processing: 4% paraformaldehyde fixation, graded sucrose dehydration, and OCT embedding. Confocal microscopy of cryosections revealed cell-specific tdTomato fluorescence distribution. Validation studies demonstrated significant spatial colocalization between endogenous Kindlin-2 (green channel) revealed by immunofluorescent (IF) staining and reporter signals (red channel) in cells located in the superficial articular cartilage (i.e., resting and proliferative articular chondrocytes) (Figure 1A-C); this result is highly consistent with that revealed by IF staining showing that Kindlin-2 is highly expressed in the articular chondrocytes^7^.

**Figure 1.**
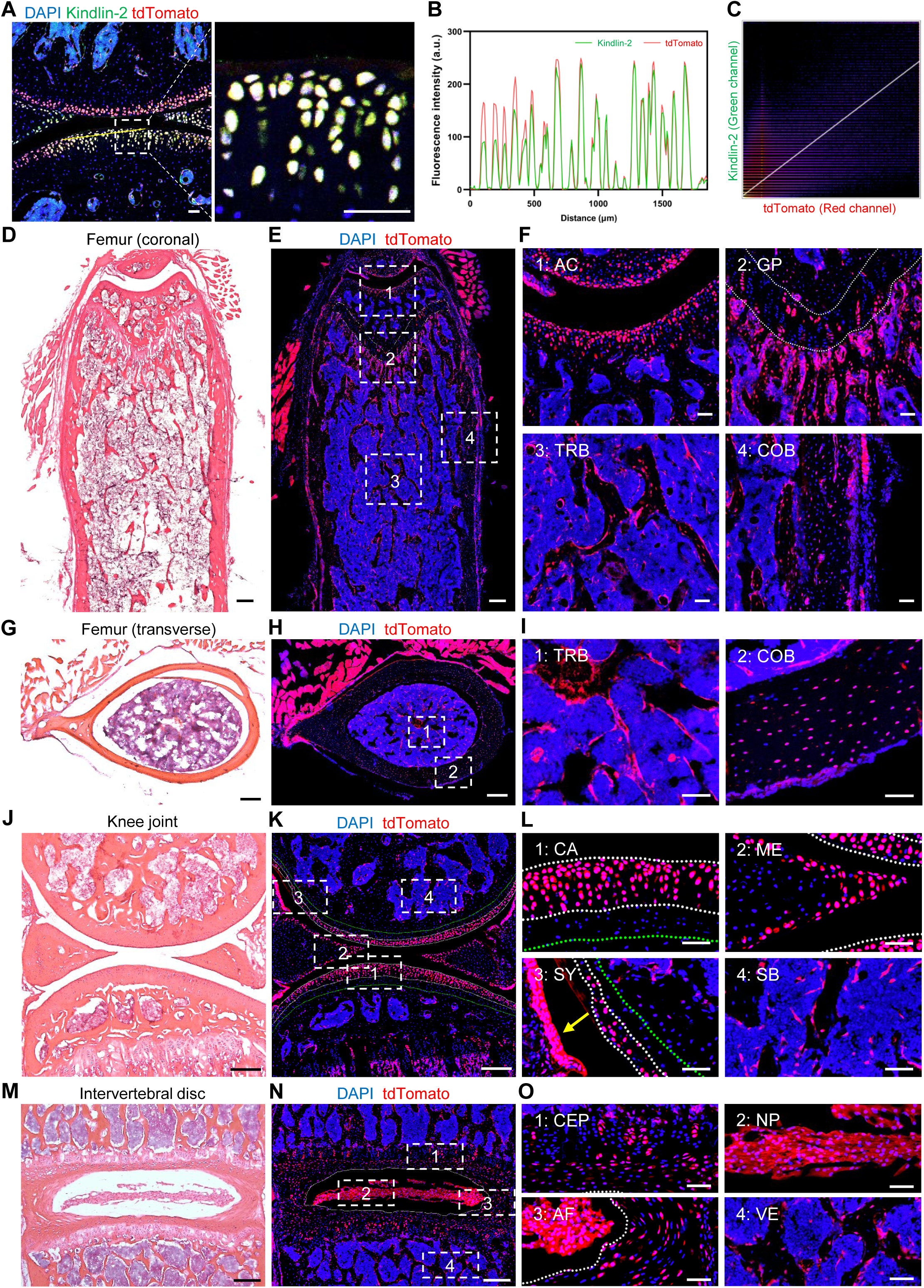
Kindlin-2 expression in murine bone and joint tissues. (A) Immunofluorescence (IF) staining of Kindlin-2 in the knee joint. White dashed rectangles indicate regions selected for magnification. (B) Correlation plot of Kindlin-2 antibody fluorescence intensity versus tdTomato fluorescence in articular cartilage. The yellow line (in A) delineated the regions subjected to analysis. (C) Colocalization analysis of Kindlin-2 antibody and tdTomato fluorescence signals in the knee joint. (D-F) H&E staining (D) and tdTomato fluorescence (E-F) in coronal sections of the femur. The growth plate region was demarcated by white dotted lines. (G-I) H&E staining (G) and tdTomato fluorescence (H-I) in transverse sections of the femur. (J-L) H&E staining (J) and tdTomato fluorescence (K-L) in sagittal sections of the knee joint. The articular cartilage surface was outlined with white dotted lines, while the calcified cartilage layer and subchondral bone region were differentiated by green dotted lines. Yellow arrow (in L) indicates the synovium. (M-O) H&E staining (M) and tdTomato fluorescence (N-O) in coronal sections of the intervertebral disc (IVD). The nucleus pulposus area was delineated by white dotted lines. Panels F, I, L and O present higher-magnification views of the boxed regions in E, H, K and N, respectively (white dashed rectangles). Scale bars, 200 μm (A, D, E, G, H, J, K, M, and N) and 50 μm (F, I, L and O). AC: articular cartilage; GP: growth plate; TRB: trabecular bone; COB: cortical bone; ME: meniscus; SY: synovium; SB: subchondral bone; CEP: cartilaginous endplate; NP: nucleus pulposus; AF: annulus fibrosus; VE: vertebra.

### Kindlin-2 expression in skeletal system

Given the well-documented preferential expression of Kindlin-2 in cartilage tissues and its critical role in skeletal formation and homeostasis, we first prioritized the skeletal systems to validate the fidelity of our tracing platform. Through analysis of *Fermt2-CreERT2 Ai9^fl/+^* mice, we observed widespread tdTomato^+^ signals in the long bones, including the articular cartilage, trabecular bone, cortical bone, and specific regions of the growth plate (Figure 1D-I and S2A-C). Notably, tdTomato signals were strongly detected in cells (mostly osteoblasts and their precursors) located on the surfaces of trabecular bones and osteocytes embedded in the cortical bone matrix (Figure S2J, K). These findings align with those by IF staining from previous reports, which have established expression of Kindlin-2 in osteocytes and its critical role in regulating bone homeostasis^8^. However, the near absence of tdTomato fluorescence within the bone marrow cavity suggests that Kindlin-2 expression is negligible in hematopoietic cells residing in this niche (Figure 1D-I and S2A-C).

Fluorescence analysis of the knee joints revealed intense tdTomato signals in superficial articular cartilage layers but not in cells located in the adjacent calcified cartilage regions (Figure 1J-L). Subsequent examinations demonstrated distinct fluorescent labeling in the meniscus surface and synovial tissues (Figure 1L). These observations corroborate the findings of Wu et al., who documented high Kindlin-2 expression in articular chondrocytes and its essential function in regulating the knee joint homeostasis^7^. A similar expression pattern was observed in small joints, including intertarsal joints of murine hindlimbs (Figure S2G-I).

Fluorescence imaging of lumbar intervertebral disc (IVD) demonstrated that Kindlin-2 was expressed in all nucleus pulposus (NP) with smaller proportions of Kindlin-2-expressing cells observed in the annulus fibrosus (AF) and cartilaginous endplates (CEP) (Figure 1M-O). These results are highly consistent with those from those by Chen et al that demonstrated Kindlin-2 expression in the IVD^9^. This interesting study demonstrated that Kindlin-2 inhibited Nlrp3 inflammasome activation in nucleus pulposus to maintain IVD homeostasis. Remarkably, the expression profile in caudal IVD mirrored that observed in lumbar IVD (Figure S2D-F).

### Kindlin-2 expression in nervous system

Current knowledge regarding Kindlin-2’s role in the nervous system remains limited, prompting our investigation into its spatial expression profile within neural tissues. Utilizing tdTomato fluorescence imaging in *Fermt2-CreERT2 Ai9^fl/+^* mice, we identified distinct expression patterns of Kindlin-2 in the brain and spinal cord. Regional analysis of brain structures revealed heterogeneous tdTomato^+^ signals, with dense punctate fluorescence observed in the olfactory bulb, neocortex, hippocampus, thalamus, and hypothalamus, while the corpus callosum white matter exhibited minimal expression (Figure 2A-C and S3C).

**Figure 2.**
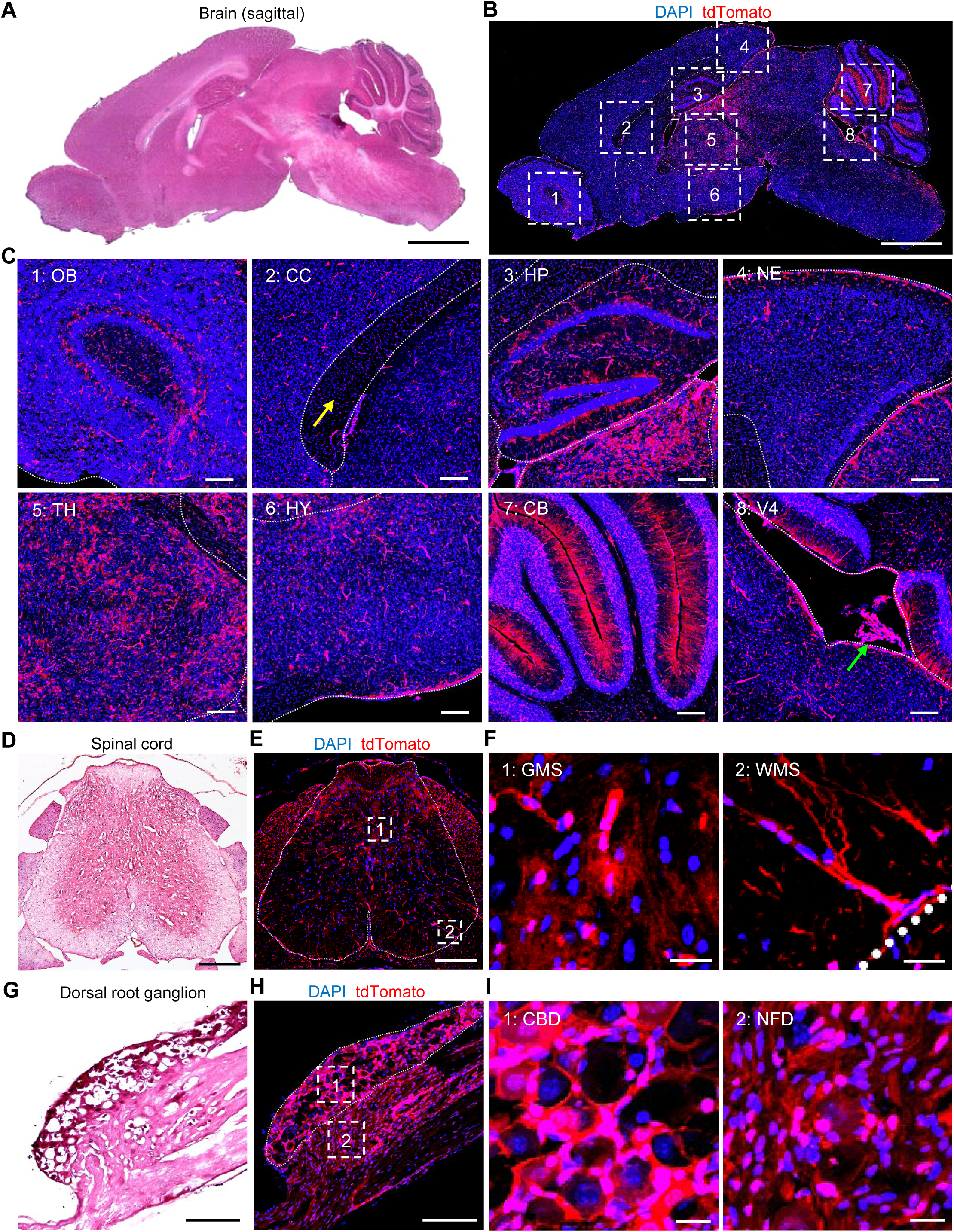
Kindlin-2 expression in the murine nervous system. (A-C) H&E staining (A) and tdTomato fluorescence (B-C) in sagittal sections of the brain. Major brain regions were demarcated by white dotted lines. Yellow arrow (in C) indicates the corpus callosum, while green arrow (in C) annotates the choroid plexus. (D-F) H&E staining (D) and tdTomato fluorescence (E-F) in transverse sections of the spinal cord. The overall spinal cord boundary was outlined with white dotted lines. (G-I) H&E staining (G) and tdTomato fluorescence (H-I) in dorsal root ganglion (DRG). Neuronal cell bodies within the DRG were delineated by white dotted lines. Panels C, F, and I show higher-magnification views of the boxed regions in B, E, and H, respectively (white dashed rectangles). Scale bars, 2000 μm (A and B), 200 μm (C, D, E, G, and H) and 20 μm (F and I). OB: olfactory bulb; CC: corpus callosum; HP: hippocampus; NE: neopallium; TH: thalamus; HY: hypothalamus; CB: cerebellum; V4: fourth ventricle; GMS: gray matter of spinal cord; WMS: white matter of spinal cord; CBD: cell bodies of dorsal root ganglia; NFD: nerve fibers of dorsal root ganglia.

Notably, the proportion of Kindlin-2 positive cells in the thalamus significantly exceeded that in the hypothalamus (Figure 2C and S3C). Cerebellar analysis demonstrated tdTomato signals in both the granular and molecular layers with high-intensity fluorescence along molecular layer fiber tracts, suggesting active Kindlin-2 transcriptional activity in these regions (Figure 2C and S3C). Previous studies have indicated Kindlin-2 expression in Purkinje neurons, suggesting partial contributions of these cells to the observed granular and molecular layer signals^10^. Additionally, distinct tdTomato fluorescence was detected in the choroid plexus of the third and fourth ventricles (Figure 2C and S3C). Colocalization analysis revealed significant overlap between tdTomato and CD31 fluorescence in cerebral vasculature and choroid plexus, confirming the vascular localization characteristics of the tdTomato signals (Figure S3D).

Spinal cord fluorescence analysis further highlighted heterogeneous Kindlin-2 expression, with tdTomato^+^ cells detected in gray matter and fiber-associated signals in white matter (Figure 2D-F). In dorsal root ganglia (DRG), Kindlin-2 localized to neuronal somata and fibers, while remaining undetectable in others (Figure 2G-I). This observation corroborates previous findings demonstrating higher Kindlin-2 concentrations in DRG compared to cerebral tissues^10^.

### Kindlin-2 expression in respiratory and circulatory system

In the murine respiratory system, we focused on lung and tracheae. Analysis of pulmonary fluorescence distribution in *Fermt2-CreERT2 Ai9^fl/+^* mice revealed a pan-pulmonary expression of Kindlin-2. tdTomato fluorescence was prominently observed in alveolar regions and the smooth muscle layer of terminal bronchioles, with specific localization to pulmonary microvasculature, while ciliated columnar epithelium in terminal bronchioles showed no detectable signals (Figure 3A-C). In contrast to bronchiolar architecture, tracheal analysis demonstrated predominant enrichment of Kindlin-2 in hyaline chondrocytes within tracheal cartilage rings (Figure 3D-F).

**Figure 3.**
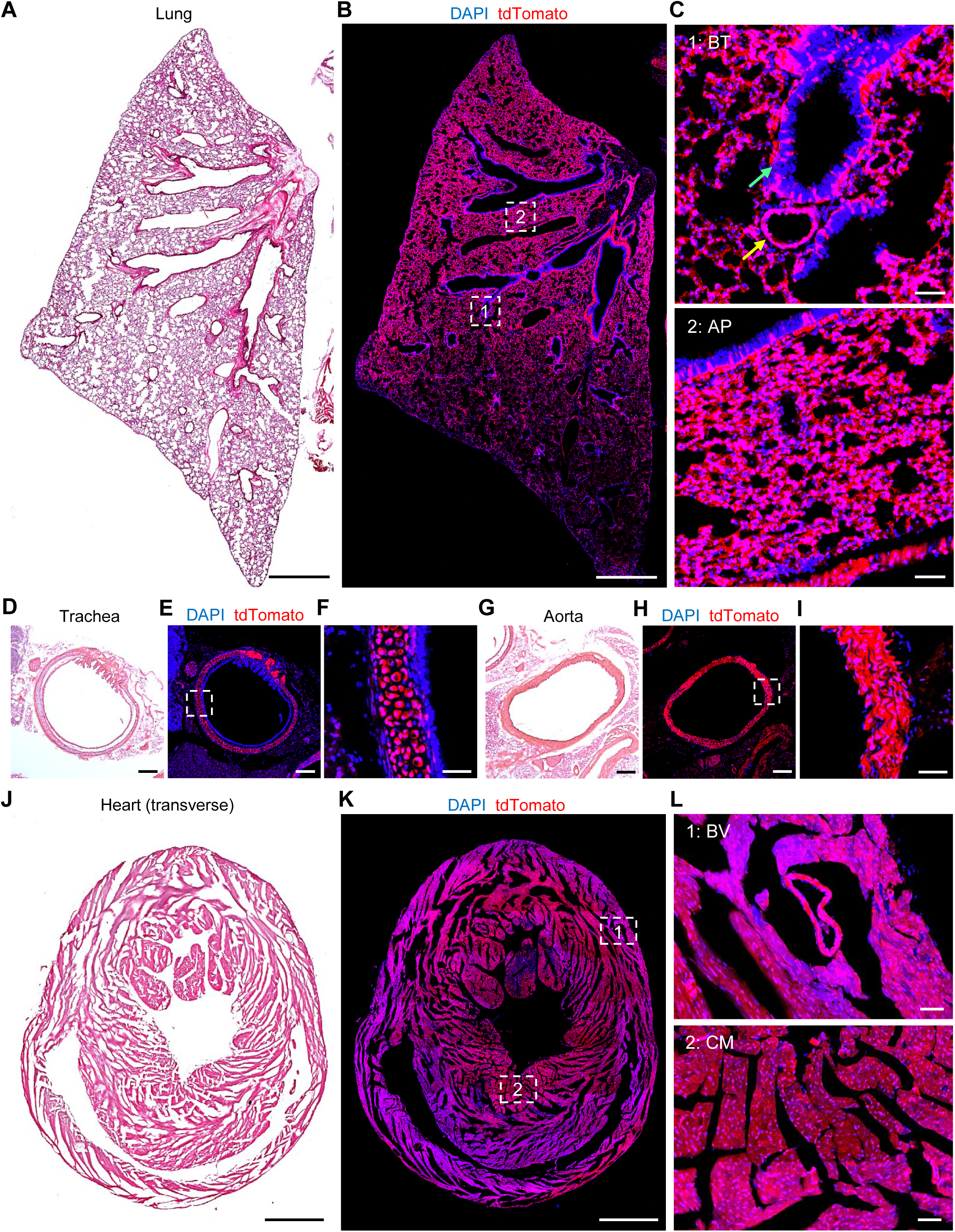
Kindlin-2 expression in murine respiratory and circulatory systems. (A-C) H&E staining (A) and tdTomato fluorescence (B-C) in maximal projection planes of the lung. Green arrow (in C) designates terminal bronchioles, while yellow arrow (in C) indicates peribronchiolar vasculature. (D-F) H&E staining (D) and tdTomato fluorescence (E-F) in transverse sections of the trachea. (G-I) H&E staining (G) and tdTomato fluorescence (H-I) in transverse sections of the aorta. (J-L) H&E staining (J) and tdTomato fluorescence (K-L) in transverse sections of the heart. Panels C, F, I, and L display higher-magnification views of the boxed regions in B, E, H, and K, respectively (white dashed rectangles). Scale bars, 1000 μm (A, B, J, and K), 200 μm (D, E, G, and H) and 50 μm (C, F, I, and L). BT: bronchioli terminals; AP: alveolus pulmonis; BV: blood vessel; CM: cardiac muscle.

Within the circulatory system, our investigation focused on cardiac and vascular structures. Analysis of tdTomato fluorescence demonstrated continuous signals in the tunica media smooth muscle layer of the aorta and inferior vena cava, while no detectable fluorescence was observed in the tunica adventitia or tunica intima (Figure 3G-I and S4D-F). Myocardial tissue exhibited strong, homogeneous tdTomato fluorescence, with cardiac vasculature also displaying Kindlin-2 positivity (Figure 3J-L and S4A-C). These findings validate previous studies reporting Kindlin-2 expression in cardiac tissues^11,12^.

### Kindlin-2 expression in digestive system

In the digestive system, we systematically characterized the spatial expression profile of Kindlin-2 across gastrointestinal (GI) tract and associated organ including liver and pancreas. Histological and tdTomato fluorescence analyses of GI tract tissues revealed no significant signals in the stratified squamous epithelium of the esophagus, with specific labeling restricted to the muscularis mucosae and muscularis propria (Figure S5A-C). Gastric wall imaging in mice similarly demonstrated predominant tdTomato fluorescence in the muscularis mucosae and muscularis propria (Figure 4A-C). Further examination of intestinal sections confirmed predominant Kindlin-2 expression in the muscularis mucosae and circular/longitudinal muscle layers of both small and large intestines, whereas no detectable expression was observed within the epithelial lining of intestinal villi (Figure 4D-F). Discrete tdTomato^+^ cell clusters were additionally detected within small intestinal Peyer’s patches (Figure 4E, F).

**Figure 4.**
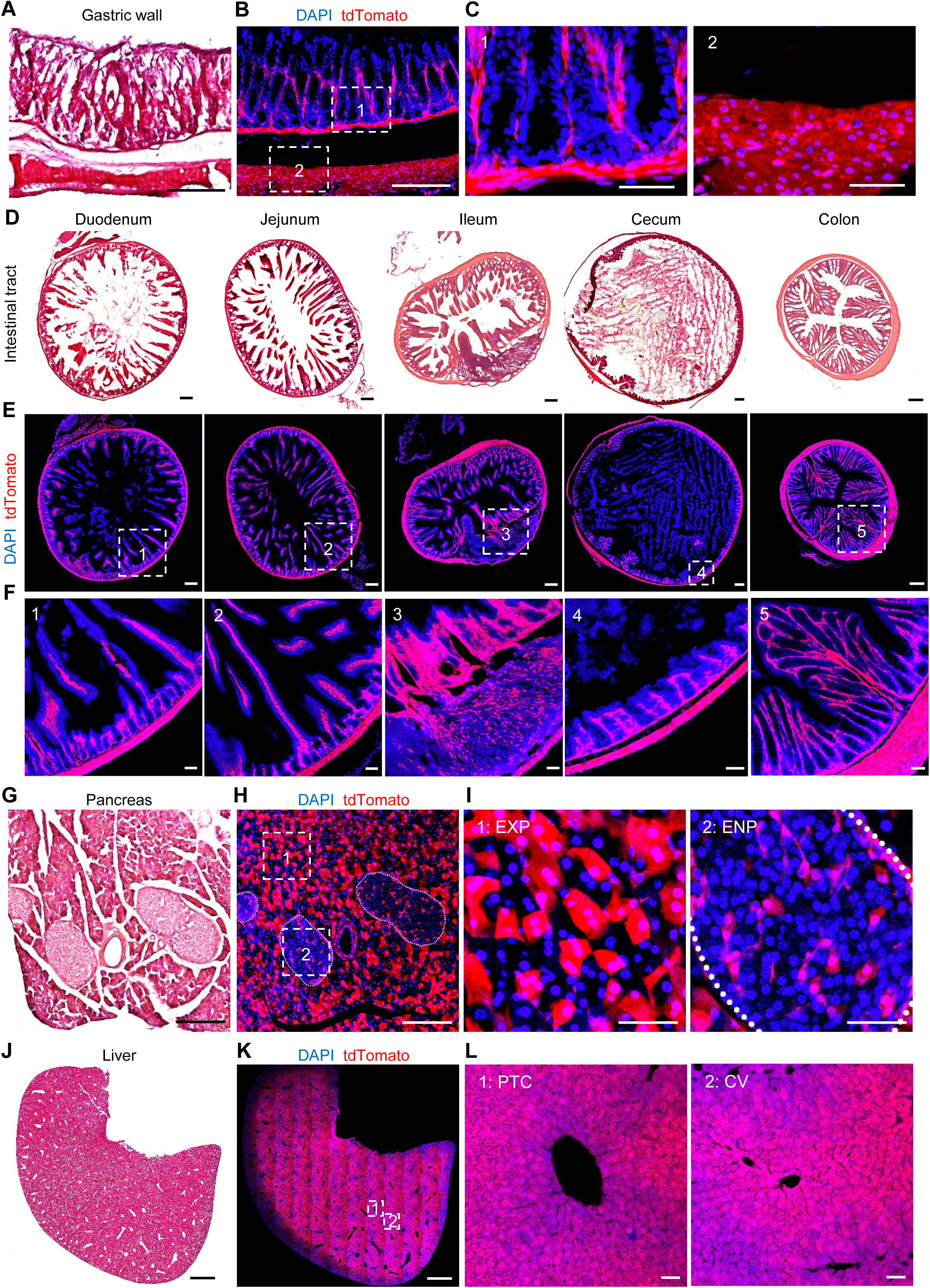
Kindlin-2 expression in the murine digestive system. (A-C) H&E staining (A) and tdTomato fluorescence (B-C) in the gastric wall. (D-F) H&E staining (D) and tdTomato fluorescence (E-F) in transverse sections of intestinal segments. (G-I) H&E staining (G) and tdTomato fluorescence (H-I) in the pancreas. Pancreas islets were demarcated by white dotted lines. (J-L) H&E staining (J) and tdTomato fluorescence (K-L) in maximal projection planes of the liver. Panels C, F, I, and L show higher-magnification views of the boxed regions in B, E, H, and K, respectively (white dashed rectangles). Scale bars, 1000 μm (J and K), 200 μm (A, B, D, E, G, and H) and 50 μm (C, F, I, and L). EXP: exocrine portion; ENP: endocrine portion; PTC: portal canal; CV: central venous.

Analysis of digestive glands revealed high Kindlin-2 expression in murine salivary glands, specifically localized to submandibular gland acinar cells, with no detectable signals in salivary ducts (Figure S5D-F). Pancreatic imaging demonstrated tdTomato fluorescence in both exocrine and endocrine compartments, though fluorescence intensity in exocrine acinar cells markedly exceeded that in islet endocrine cells (Figure 4G-I). Gallbladder analysis showed Kindlin-2 expression confined to the smooth muscle layer, with no detection in the columnar epithelial lining (Figure S5G-I). Notably, hepatic tissue exhibited intense tdTomato signals throughout the parenchyma, with perivenous hepatocytes demonstrating stronger fluorescence compared to periportal regions (Figure 4J-L).

### Kindlin-2 expression in urinary and reproductive system

In the urinary system, Kindlin-2 exhibited marked regional specificity. Renal cortical analysis revealed tdTomato fluorescence predominantly localized to scattered glomeruli, with minimal signal detected across cortical tubules. (Figure 5A-C). In contrast, intense Kindlin-2 positivity was observed in the renal medulla, particularly within the loop of Henle and renal papilla. Additional tdTomato signals were detected surrounding convoluted tubules and collecting ducts (Figure 5A-C). Adrenal glands also displayed tdTomato fluorescence (Figure 5C). Bladder analysis demonstrated Kindlin-2 expression restricted to the muscularis mucosae and smooth muscle layers, with no detectable signals in the mucosal columnar epithelium (Figure 5D-F). Similarly, murine penile tissues showed absence of tdTomato fluorescence in urethral epithelium but revealed numerous Kindlin-2 positive cells within the corpus cavernosum and vasculature (Figure 5G-I).

**Figure 5.**
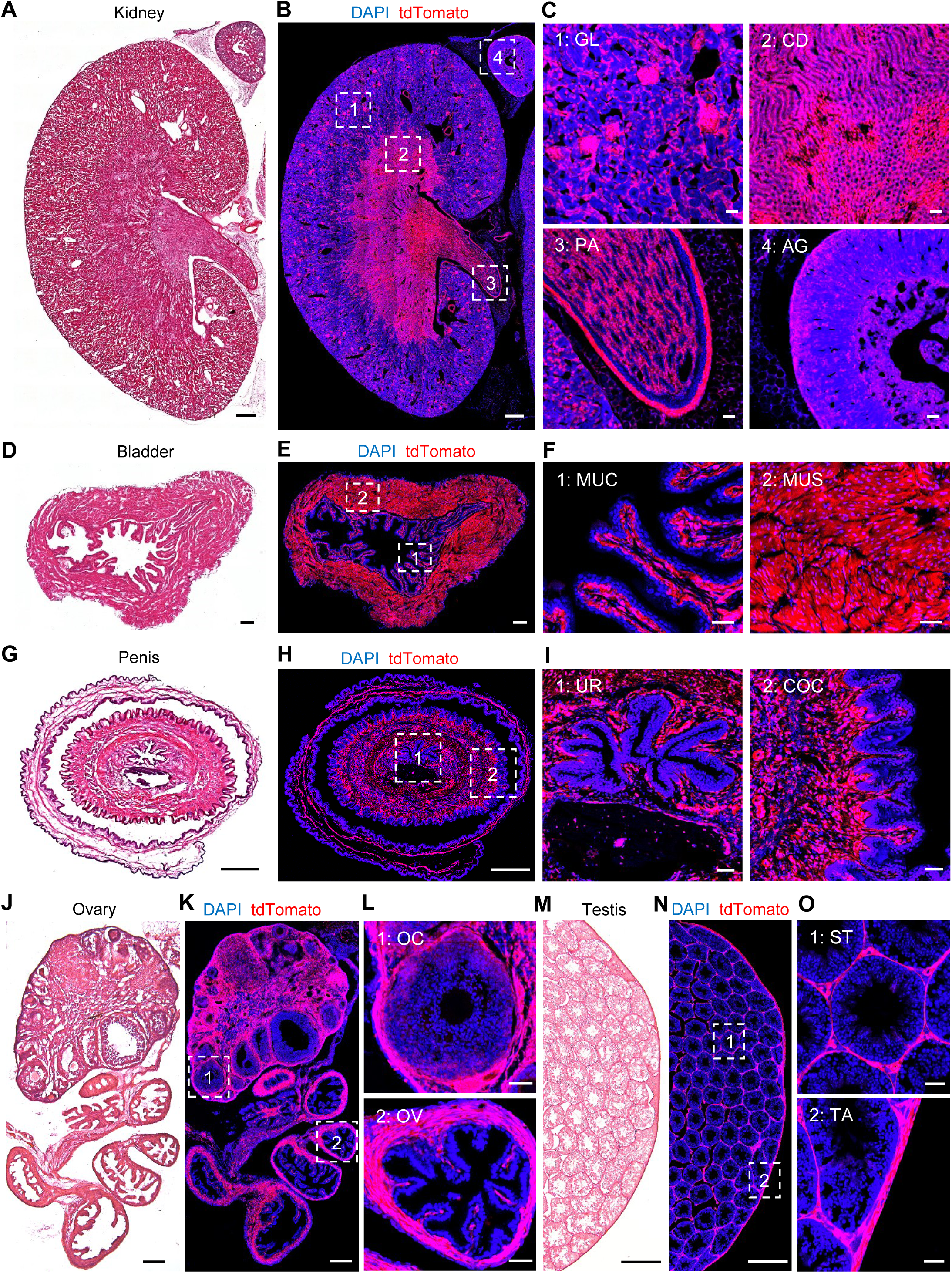
Kindlin-2 expression in the murine urogenital system. (A-C) H&E staining (A) and tdTomato fluorescence (B-C) in coronal sections of the kidney. (D-F) H&E staining (D) and tdTomato fluorescence (E-F) in the bladder. (G-I) H&E staining (G) and tdTomato fluorescence (H-I) in transverse sections of the penis. (J-L) H&E staining (J) and tdTomato fluorescence (K-L) in the ovary and oviduct. (M-O) H&E staining (M) and tdTomato fluorescence (N-O) in transverse sections of the testis (partial). Panels C, F, I, L, and O display higher-magnification views of the boxed regions in B, E, H, K, and N, respectively (white dashed rectangles). Scale bars, 500 μm (A, B, G, H, M and N), 200 μm (D, E, J, and K) and 50 μm (C, F, I, L, and O). GL: glomerulus; CD: collecting duct; PA: renal papillae; AG: adrenal gland; MUC: mucosa; MUS: muscularis; UR: urethra; COC: corpus cavernosum; OC: oocyte; OV: oviduct; ST: seminiferous tubule; TA: tunica albuginea.

Within the reproductive system, ovarian and testicular sections from female and male mice, respectively, were analyzed. Ovarian tdTomato fluorescence localized exclusively to non-germ cells, with expression observed in the highly vascularized central medulla, and the compact outer cortex comprising developing follicles and corpora lutea (Figure 5J-L). The oviduct, sharing structural similarities with intestinal tissue, exhibited tdTomato signals in its smooth muscle layer (Figure 5L). Intriguingly, testicular expression patterns mirrored ovarian observations, showing no germ cell involvement. Fluorescence predominated in the tunica albuginea and basal lamina enveloping seminiferous tubules (Figure 5M-O). Scattered tdTomato signals were additionally detected in Leydig cells and vasculature (Figure 5O). Epididymal analysis further confirmed Kindlin-2 expression in the smooth muscle walls of both caput and cauda epididymis (Figure S6A-C).

### Kindlin-2 expression in immune system

Within the immune system, we analyzed lymphoid organs, including the thymus, spleen, and lymph nodes. Thymic analysis revealed tdTomato fluorescence predominantly localized to the medulla and vascular walls, with sparse distribution in the cortex and thymic capsule (Figure 6A-C). Splenic examination demonstrated Kindlin-2-positive cells distributed across both white pulp and red pulp, with higher fluorescence density observed in the red pulp (Figure 6D-F). Additionally, tdTomato signals were detected in central arteries and the splenic capsule, the latter likely originating from smooth muscle fibers (Figure 6F). Lymph node profiling identified minimal fluorescence in the superficial cortex and capsule. tdTomato signals were predominantly concentrated in the paracortex and medulla (Figure 6G-I).

**Figure 6.**
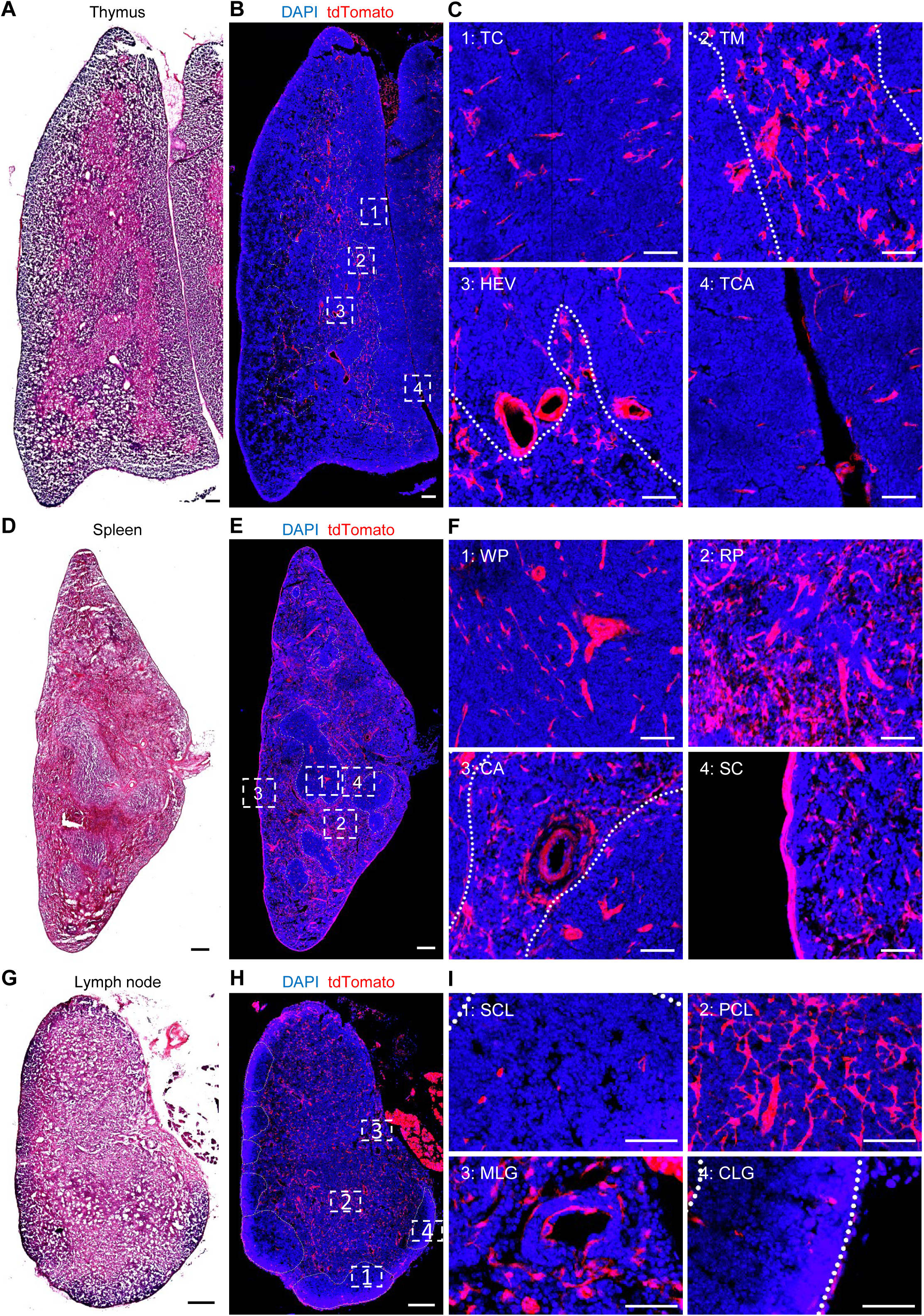
Kindlin-2 expression in the murine immune system. (A-C) H&E staining (A) and tdTomato fluorescence (B-C) in maximal projection planes of the thymus. The cortical-medullary junction was demarcated by white dotted lines. (D-F) H&E staining (D) and tdTomato fluorescence (E-F) in transverse sections of the spleen. Red pulp and white pulp regions were differentiated by white dotted lines. (G-I) H&E staining (G) and tdTomato fluorescence (H-I) in lymph node. The superficial cortex of the lymph node was outlined with white dotted lines. Panels C, F, and I present higher-magnification views of the boxed regions in B, E, and H, respectively (white dashed rectangles). Scale bars, 200 μm (A, B, D, E, G, and H) and 50 μm (C, F, and I). TC: thymic cortex; TM: thymic medulla; HEV: highly endothelial venule; TCA: thymic capsule; WP: white pulp; RP: red pulp; CA: central artery; SC: splenic capsule; SCL: superficial cortex of lymph gland; PCL: paracortex of lymph gland; MLG: medulla of lymph gland; CLG: capsule of lymph gland.

### Kindlin-2 expression in eye, adipose and skin tissues

Beyond major organ systems, we further identified Kindlin-2 expression in specialized anatomical structures. Ocular analyses revealed sparse but distinct tdTomato signals in choroid layers, retinal ganglion/inner nuclear layers, lens epithelia, ciliary bodies, and corneal stroma (Figure 7A-C). Adipose tissue examinations demonstrated abundant tdTomato fluorescence in both brown and white adipose depots (Figure 7D-G). Murine dorsal skin analysis showed no detectable Kindlin-2 signals in the epidermal layer (Figure 7H-J). Within the dermis, tdTomato fluorescence was observed in arrector pili muscles and vasculature, with notable signals in hair follicles warranting further investigation (Figure 7J and S7A-C). Subcutaneous muscular layers exhibited the most intense fluorescence in skin tissues (Figure 7J). Significant tdTomato signals were similarly detected in skeletal muscles (Figure S7D, E) and tenocytes (Figure S7F-H).

**Figure 7.**
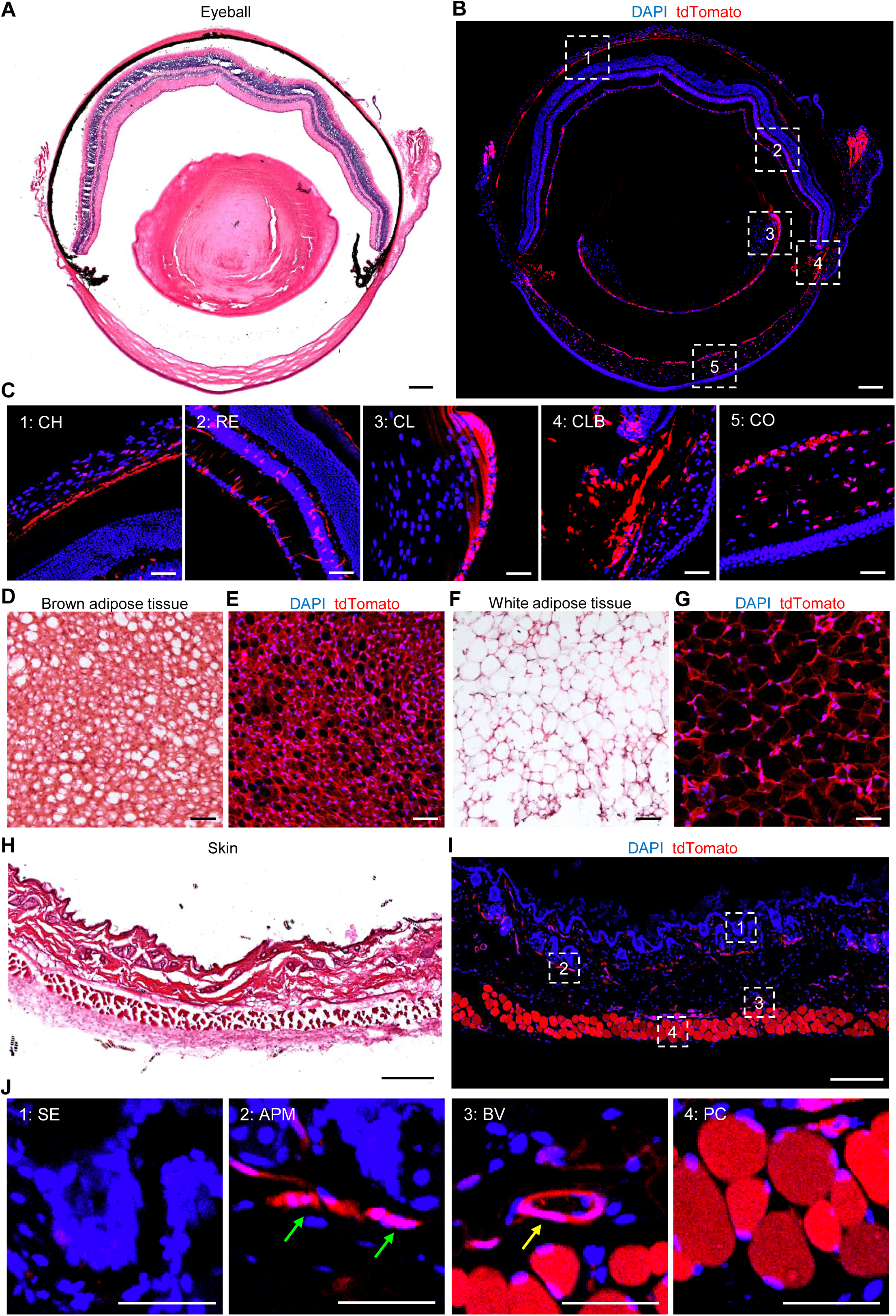
Kindlin-2 expression in eye, adipose and skin tissues. (A-C) H&E staining (A) and tdTomato fluorescence (B-C) in transverse sections of the eyeball. (D-E) H&E staining (D) and tdTomato fluorescence (E) in brown adipose tissue (BAT). (F-G) H&E staining (F) and tdTomato fluorescence (G) in white adipose tissue (WAT). (H-J) H&E staining (H) and tdTomato fluorescence (I-J) in skin. In J, the arrector pili muscle is denoted by green arrows, while subcutaneous vasculature is indicated by yellow arrow. Panels C and J show higher-magnification views of the boxed regions in B and I, respectively (white dashed rectangles). Scale bars, 200 μm (A, B, H, and I) and 50 μm (C, D, E, F, G and J). CH: choroid; RE: retina; CL: crystalline lens; CLB: ciliary body; CO: cornea; SE: surface epithelium; APM: arrector pili muscle; PC: panniculus carnosus.

## DISCUSSION

The CreERT2/Ai9 system combined with a “so-called” tissue-specific promoter deriving Cre recombinase expression has been employed to delineate spatiotemporal expression patterns of regulators^6^. However, its application for organism-wide spatial expression analysis remains virtually unexplored. In this study, for the first time to our knowledge, we have engineered an inducible *Fermt2-CreERT2* knock-in mouse model, enabling systemic interrogation of Kindlin-2’s pan-tissue/organ expression patterns across temporally restricted phases. The inducible X-CreERT2/Ai9 system utilized in this study can directly couple fluorescent output to transcriptional activity of the target gene promoter, ensuring superior specificity and stability. It demonstrates superior performance to conventional detection methodologies in terms of fidelity and resolution, offering significant advantages for profiling target gene expression within defined temporal windows.

Importantly, the Kindlin-2 expression patterns revealed by the *Fermt2-CreERT2/Ai9* system in this study are highly consistent with those published in published literatures. For example, in skeleton, Kindlin-2 expression has been reported to be expressed in osteocytes, osteoblasts, chondrocytes, and nucleus pulposus cells^7–9,13^, which aligns with results from this study. Similarly, in the circulatory system, existing studies localize Kindlin-2 to cardiac Z-discs and vascular smooth muscle^11,12,14–19^, findings that are highly consistent with results from this study. These results demonstrate the high-fidelity of the novel tracing system from this study.

Although we have detected Kindlin-2 expression in multiple neural-associated tissues, its roles in the nervous system remain largely underexplored, with only limited studies linking it to neuronal axon elongation^10,20^. Within the respiratory system, prior research has primarily reported Kindlin-2 expression in the lung^21–25^, while our study additionally identifies robust expression in tracheal hyaline cartilage and its complete absence in ciliated columnar epithelium. The digestive system exhibits predominant Kindlin-2 expression in the gastrointestinal tract, liver, and pancreas^19,26–30^. In the urinary system, Kindlin-2 has been implicated in renal podocytes and penile tissues, as reported previously^31–36^. Our work not only corroborates these observations but also reveals broader renal expression domains. In this study, we identify Kindlin-2 expression in thymic, splenic, and lymph node tissues. Furthermore, we validate its expression in adipose tissue^37–39^. While these findings further exhibit strong concordance with established literature, we demonstrate previously undocumented Kindlin-2 expression in the brain, spinal cord, DRG, trachea, thymus, spleen, lymph nodes, and ocular structures. These discoveries address critical gaps in understanding Kindlin-2’s pathophysiological roles and establish a foundational atlas for future functional studies in understudied cells and tissues.

It is also important that we have identified distinct anatomical compartments exhibiting near-undetectable Kindlin-2 expression. These include mucosal epithelia (gastrointestinal, respiratory, and urinary tracts), germ cells (oocytes and spermatogenic cells), avascular tubular structures (renal cortical tubules and salivary gland ducts), and the epidermal layer of the skin. This absence suggests that Kindlin-2 does not play a direct role in cells of these anatomical compartments.

It should be noted that there are marked discrepancies between the Kindlin-2 expression patterns in pancreas and testicle revealed by the *Fermt2-CreERT2/Ai9* system in this study and by IF staining in literature. Zhu et al. previously localized Kindlin-2 predominantly to pancreatic β-cells using IF^28^. In contrast, results from this study reveal dual expression in both exocrine acinar cells and islets, with markedly enhanced tdTomato fluorescence intensity in exocrine compartments. This divergence may stem from technical limitations in antibody-based detection; the dense cytoplasmic architecture of acinar cells likely impeded antibody permeabilization, preventing epitope recognition in prior methodologies. Similarly, our testicular analyses conflict with established reports. While existing studies implicate Kindlin-2 expression in Sertoli and germ cells^40^, our genetic tracing identifies predominant expression within the seminiferous tubule walls, with no detectable signals in Sertoli or germline populations. This spatial discordance warrants further investigation in the future.

Contemporary multi-omics research predominantly employs transcriptomic and proteomic sequencing technologies, which utilize high-throughput analytical strategies for parallel gene detection, constructing molecular expression profiles across biological samples in a horizontal dimension. In contrast, the inducible X-CreERT2/Ai9 system adopts a divergent approach, enabling longitudinal tracking of single-gene expression through precise spatial control. This methodology facilitates in-depth functional dissection across developmental trajectories and pathological progression in model organisms. Simultaneous integration of the X-CreERT2/Ai9 system with multi-omics methodologies enables precise histological classification of target gene-expressing cells, thereby elucidating the regulatory principles governing their expression dynamics. Notably, the integration of TAM-inducible Cre recombinase establishes a regulated gene expression platform. This innovation transcends the static detection limitations of conventional omics by permitting targeted gene activation at defined developmental stages or disease states, thereby dynamically resolving temporal-spatial correlations between expression patterns and pathophysiological processes. Furthermore, future investigations could leverage artificial intelligence algorithms integrated with volumetric tissue imaging to reconstruct three-dimensional expression atlases, thereby deciphering target gene expression signatures at higher-dimensional resolution.

In pharmacological target validation, the system may demonstrate its unique applications in the future. Despite growing emphasis on targeted therapies, critical knowledge gaps persist regarding the dynamic expression profiles of therapeutic targets and their interaction with drug candidates. The X-CreERT2/Ai9 system addresses this by enabling precise tracing of target gene expression, offering a novel paradigm to delineate therapeutic windows, optimize dosing regimens, and deconvolve drug resistance mechanisms. Current research on drug targets remains superficial, exploring additional expression loci may unveil unanticipated functionalities, potentially reshaping existing biological paradigms. Crucially, the platform’s modular architecture confers functional extensibility. Through incorporation of molecular adaptors between target genes and CreERT2, this methodology extends to expressive studies of non-coding genomic elements (e.g., microRNAs), providing technical support for dynamic interrogation of epigenetic regulatory networks. Such versatility not only advances mechanistic understanding of gene regulation but also positions the system as an innovative toolkit for personalized therapeutic design in precision medicine.

### Limitations of the study

While the X-CreERT2/Ai9 system provides robust spatiotemporal resolution for gene expression profiling, its inherent limitations necessitate critical consideration. First, a potential spatial segregation between the target protein and the cytoplasmic tdTomato reporter precludes precise subcellular localization analysis. Although fusion constructs of the target protein with fluorescent tags could resolve this, such modifications risk altering native protein functionality. Second, the fluorescence intensity reflects cumulative tdTomato expression over time post-TAM induction rather than real-time target gene expression levels, introducing temporal confounding in quantitative assessments. Semi-quantitative detection of real-time target gene expression may be achieved via IF staining. Third, despite the reporter’s stability, tdTomato remains susceptible to photobleaching under suboptimal conditions (e.g., elevated temperatures or pH fluctuations), requiring stringent experimental controls during tissue processing. Photobleaching can be mitigated through low-temperature and light-restricted protocols. Fourth, the requirement for genetic manipulation restricts its application to small animal models, precluding studies in humans. However, transgenic models in large animals, such as non-human primates, may bridge this gap by recapitulating human-like gene expression profiles.

## STAR**★**METHODS

Detailed methods are provided in the online version of this paper and include the following:

- KEY RESOURCES TABLE
- RESOURCE AVAILABILITY

`o` Lead contact
`o` Materials availability
`o` Data and code availability
- EXPERIMENTAL MODEL AND SUBJECT DETAILS
- METHOD DETAILS

`o` Sample Preparation
`o` Cryosectioning Protocol
`o` Histological Staining
`o` Immunofluorescence Staining
- QUANTIFICATION AND STATISTICAL ANALYSIS

## ACKNOWLEDGMENTS

The authors acknowledge the assistance of Core Research Facilities of Southern University of Science and Technology. This work was supported, in part, the Shenzhen Medical Research Funds (B2402033), the Shenzhen Fundamental Research Program (JCYJ20220818100617036), the National Natural Science Foundation of China Grants (82430078, 82230081, 82261160395, 82250710175), the Guangdong Provincial Science and Technology Innovation Council Grant (2017B030301018), and the Shenzhen Key Laboratory of Cell Microenvironment Grant (ZDSYS20140509142721429).

## AUTHOR CONTRIBUTIONS

Study design: GX and FL. Study conduct and data collection: FL. Data analysis: GX and FL. Data interpretation: GX and FL. Drafting the manuscript: GX and FL. GX and FL take the responsibility for the integrity of the data analysis.

## DECLARATION OF INTERESTS

The authors declare that they have no competing financial interests.

## STAR★METHODS

KEY RESOURCES TABLE

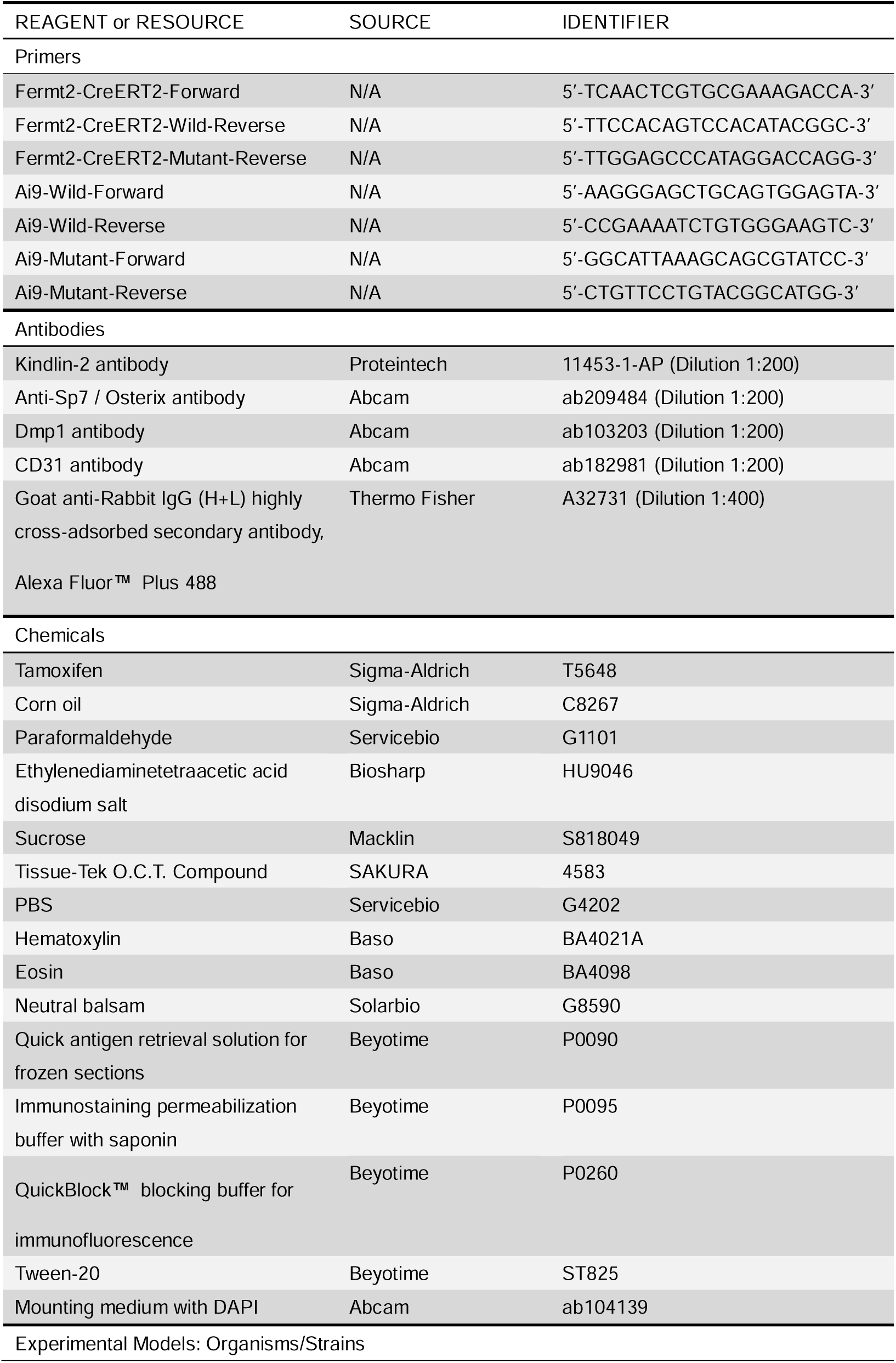
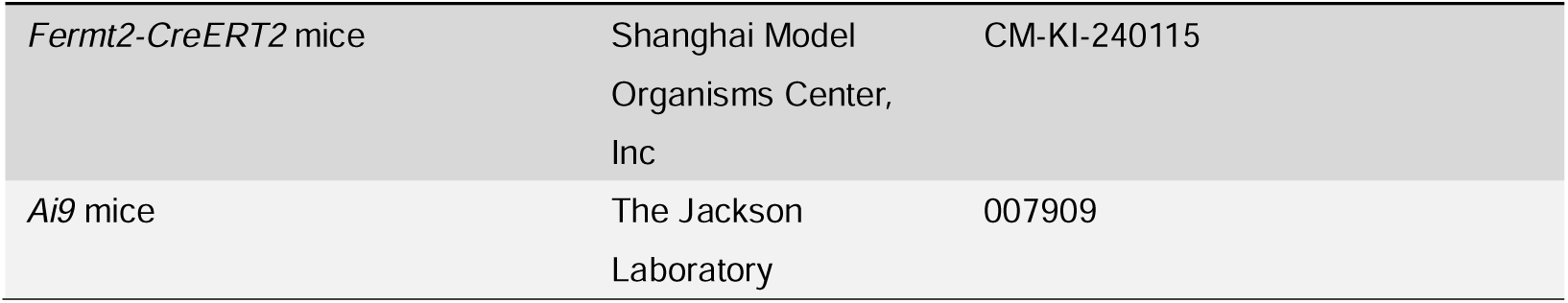

## RESOURCE AVAILABILITY

### Lead contact

Further information and requests should be directed to and will be fulfilled by the lead contact, Guozhi Xiao.

### Materials availability

All materials employed in the present investigation were sourced from commercial suppliers as specified in the Key Resources Table.

### Data and code availability

The complete set of original image data presented in this study is accessible from the corresponding author on request.

## EXPERIMENTAL MODEL AND SUBJECT DETAILS

All experimental protocols involving mice in this study were approved by the Institutional Animal Care and Use Committee (IACUC) of Southern University of Science and Technology (Approval No: SUSTech-JY202503128). Mice were maintained on a C57BL/6J genetic background under standardized housing conditions with a 12-hour light/dark cycle and ambient temperature of 20–24°C. The *Fermt2-CreERT2* strain (*C57BL/6J-Fermt2-em1(2A-CreERT2)Smoc*) was custom-generated via CRISPR/Cas9-mediated targeting by Shanghai Model Organisms Center, Inc. *Ai9* reporter mice

(*B6.Cg-Gt(ROSA)26Sor-tm9(CAG-tdTomato)Hze/J*) were derived from The Jackson Laboratory. Genetic crosses between heterozygous *Fermt2-CreERT2* and homozygous *Ai9* mice yielded *Fermt2-CreERT2 Ai9^fl/+^* offspring for lineage tracing.

## METHOD DETAILS

### Sample Preparation

Neonatal mice were genotyped by PCR to identify *Fermt2-CreERT2 Ai9^fl/+^* littermates, using primers listed in the Key Resources Table. Two-month-old *Fermt2-CreERT2 Ai9^fl/+^* mice received daily intraperitoneal injections of TAM (100 mg/kg body weight; Sigma-Aldrich, T5648) for 5 consecutive days, while genetically matched controls were administered equivalent volumes of corn oil (Sigma-Aldrich, C8267) under identical protocols. Thirty days post-induction, multiple tissues were harvested and fixed in 4% paraformaldehyde (PFA; Servicebio, G1101) for 24 hours. Bone and joint specimens underwent additional decalcification in ethylenediaminetetraacetic acid disodium salt (EDTA · 2Na; Biosharp, HU9046) for 21 days following fixation.

### Cryosectioning Protocol

Following sample preparation, tissues underwent cryoprotection through graded sucrose infiltration: initial immersion in 20% sucrose (Macklin, S818049) until gravitational sedimentation, followed by 30% sucrose for 48 hours.

Specimens were then sequentially equilibrated in Optimal Cutting Temperature (O.C.T.) compound (SAKURA, 4583) using a three-stage protocol: (1) 1:1 (v/v) 20% sucrose/OCT mixture for 2 hours at room temperature (RT, 22±3°C), (2) immersion in pure O.C.T. for 4 hours, followed by (3) fresh O.C.T. incubation for 6 hours. Cryosectioning was performed using a Leica CM1950 cryostat maintained at -25°C, generating 20 μm-thick sections that were mounted on pre-coated slides and archived at -20°C for subsequent analyses.

### Histological Staining

Cryosections were equilibrated in 1×phosphate buffered saline (PBS; Servicebio, G4202) for 30 minutes at RT to remove residual O.C.T. compound. Hematoxylin staining (Baso, BA4021A) was performed for 2-3 minutes followed by three aqueous rinses. Sections were differentiated in acid alcohol (1% HCl in 70% ethanol), counterstained 10-20 seconds with eosin (Baso, BA4098), and rinsed sequentially. Dehydration was achieved through a graded ethanol series (75%, 95%, 100%) prior to xylene clearing. Mounting with neutral resin (Solarbio, G8590) preceded whole-slide digitization using a Leica Aperio VERSA 8 scanning platform at 10× resolution.

### Immunofluorescence Staining

Cryosections were equilibrated in 1×PBS for 30 minutes at RT to remove residual O.C.T. compound. Antigen retrieval was performed using frozen section antigen recovery solution (Beyotime, P0090), followed by membrane permeabilization with immunostaining permeabilization buffer (Beyotime, P0095). Non-specific binding was blocked using immunostaining blocking buffer (Beyotime, P0260). Primary antibodies (1:200) incubation proceeded at 4°C for 16 hours in a humidified chamber. Unbound antibodies were removed through three 5-minute PBST washes (1×PBS + 0.1% Tween-20; Beyotime, ST825). Alexa Fluor 488-conjugated secondary antibodies (1:400) were applied at RT for 1 hour, followed by DAPI nuclear counterstaining using antifade mounting medium (Abcam, ab104139). Confocal imaging was conducted on a Zeiss LSM 980 system. All antibody specifications are provided in the Key Resources Table.

## QUANTIFICATION AND STATISTICAL ANALYSIS

Colocalization quantification was performed using ImageJ software (version 1.54p) with the Coloc 2 plugin.

## Supplementary figure legends

**Figure S1. Generation of the *Fermt2-CreERT2/Ai9* reporter system.** (A) Schematic diagram of core sequences in the *Fermt2-CreERT2* and *Ai9* reporter mice. (B) Operational mechanism of the *Fermt2-CreERT2/Ai9* lineage-tracing system. (C) Experimental timeline for transgenic mouse studies. (D) Gross morphology of 3-month-old control and TAM-induced mice. Scale bars, 1 mm (D).

**Figure S2. Kindlin-2 expression in additional murine bone and joint tissues, related to Figure 1**.

(A-C) H&E staining (A) and tdTomato fluorescence (B-C) in coronal sections of the tibia. (D-F) H&E staining (D) and tdTomato fluorescence (E-F) in caudal IVD. (G-I) H&E staining (G) and tdTomato fluorescence (H-I) in sagittal sections of the paw. (J) IF staining of Osterix in femoral trabecular bone. White dashed rectangles denote regions selected for magnification. Yellow arrows indicate cells exhibiting colocalization of Osterix (green fluorescence) and Tomato signals. (K) IF staining of Dmp1 in femoral cortical bone. White dashed rectangles mark magnified regions. Yellow arrows highlight cells with overlapping Dmp1 (green fluorescence) and Tomato signals. Panels C, F, and I display higher-magnification views of the boxed regions in B, E, and H, respectively (white dashed rectangles). Scale bars, 1000 μm (G and H), 200 μm (A, B, D, E, and I) and 50 μm (C, F, J and K). IJ: intertarsal joints; FP: foot pad.

**Figure S3. Expression and localization of Kindlin-2 in murine brain tissue, related to Figure 2.**

(A-C) H&E staining (A) and tdTomato fluorescence (B-C) in coronal sections of the brain. In C, green arrow (in C) denotes the olfactory bulb limb, yellow arrow (in C) indicates the subependymal zone of the lateral ventricle, and white arrows mark the choroid plexus. (D) IF staining of CD31 in the murine brain. Panel C presents a higher-magnification view of the boxed regions in B (white dashed rectangles). Scale bars, 1000 μm (A and B) and 50 μm (C and D). ONL: olfactory nerve layer; MOB: main olfactory bulb; AON: anterior olfactory nucleus; LV: lateral ventricle; CP: choroid plexus.

**Figure S4. Expression of Kindlin-2 in murine heart and veins, related to Figure 3.**

(A-C) H&E staining (A) and tdTomato fluorescence (B-C) in longitudinal sections of the heart. (D-F) H&E staining (D) and tdTomato fluorescence (E-F) in longitudinal sections of the inferior vena cava (IVC). Panels C and F display higher-magnification views of the boxed regions in B and E, respectively (white dashed rectangles). In F, vascular wall layers are annotated: tunica intima (white arrow), tunica media (yellow arrow), and tunica externa (green arrow). Scale bars, 1000 μm (A and B) and 50 μm (C, D, E, and F).

**Figure S5. Expression of Kindlin-2 in murine digestive structures, related to Figure 4.**

(A-C) H&E staining (A) and tdTomato fluorescence (B-C) in transverse sections of the esophagus. (D-F) H&E staining (D) and tdTomato fluorescence (E-F) in the salivary glands. (G-I) H&E staining (G) and tdTomato fluorescence (H-I) in the gallbladder. Panels C, F, and I show higher-magnification views of the boxed regions in B, E, and H, respectively (white dashed rectangles). Scale bars, 1000 μm (D and E), 200 μm (A, B, G, and H) and 50 μm (C, F, and I). SL: sublingual gland; SM: submandibular gland.

**Figure S6. Expression of Kindlin-2 in murine reproductive structures, related to Figure 5.**

(A-C) H&E staining (A) and tdTomato fluorescence (B-C) in transverse sections of the epididymis. Panel C presents a higher-magnification view of the boxed regions in B (white dashed rectangles). Scale bars, 200 μm (A and B) and 50 μm (C).

**Figure S7. Expression of Kindlin-2 in additional murine structures, related to Figure 7.**

(A-C) H&E staining (A) and tdTomato fluorescence (B-C) in the hair follicles of the skin. (D-E) H&E staining (D) and tdTomato fluorescence (E) in gastrocnemius. (F-H) H&E staining (F) and tdTomato fluorescence (G-H) in tendon. Panels C and H display higher-magnification views of the boxed regions in B and G, respectively (white dashed rectangles). Scale bars, 200 μm (A, B, F, and G) and 50 μm (C, D, E, and H).

## Notes

### Competing Interest Statement

The authors have declared no competing interest.

