## Supplemental Figure1-7; Supplemental Table1 for "Organism-wide spatiotemporal profiling of gene expression utilizing a X-CreERT2/Ai9 tracing system"

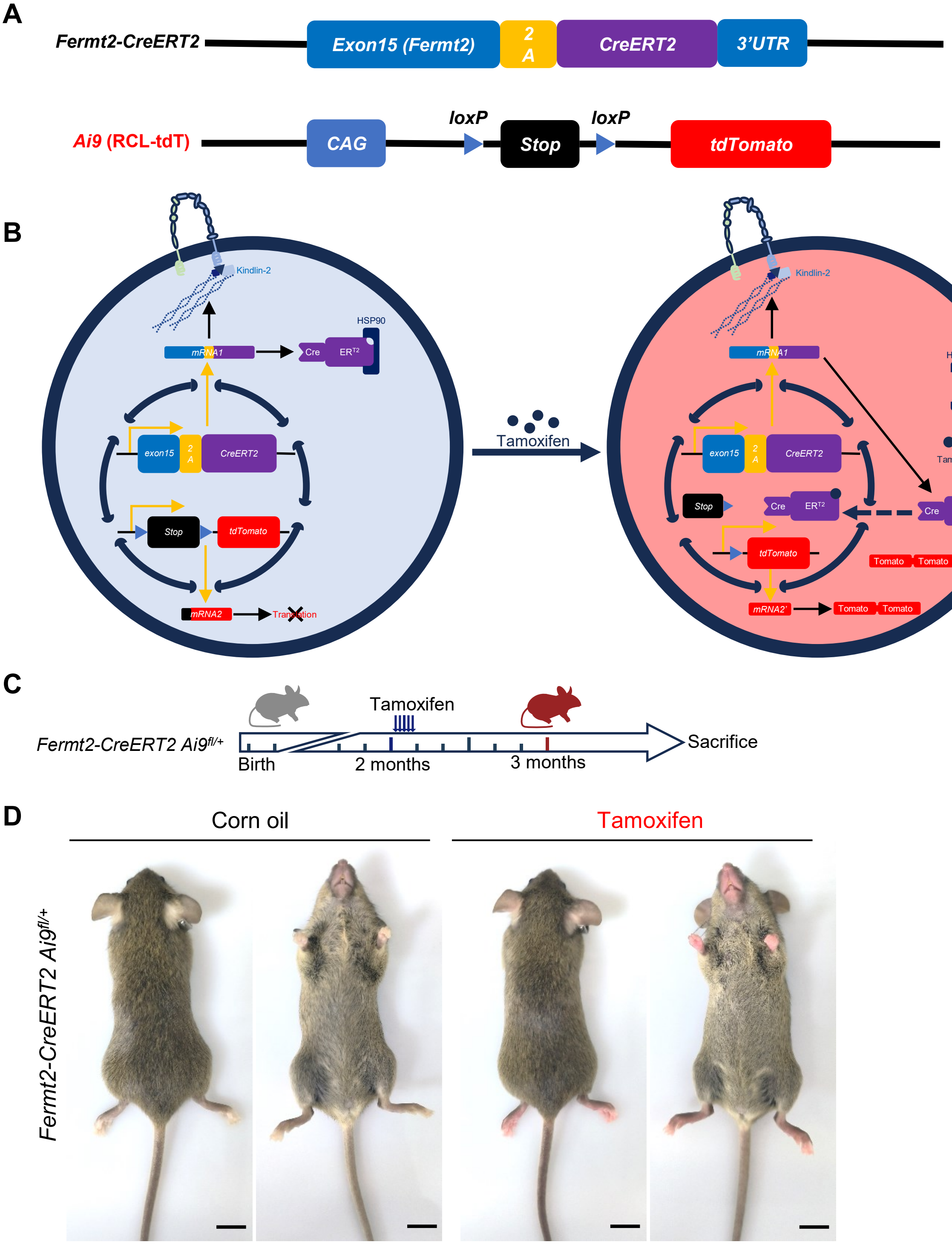
A
Fermt2-CreERT2
Exon15 (Fermt2)
2 A
CreERT2
3'UTR
Ai9 (RCL-tdT)
CAG
loxP
Stop
loxP
tdTomato
B
Kindlin-2
HSP90
mRNA1
Cre
ER T2
exon15
2 A
CreERT2
Stop
tdTomato
mRNA2
Translation
Tamoxifen
mRNA2'
Tomato
C
Fermt2-CreERT2 Ai9fl/+
Birth
Tamoxifen
2 months
3 months
Sacrifice
D
Corn oil
Tamoxifen
Fermt2-CreERT2 Ai9fl/+

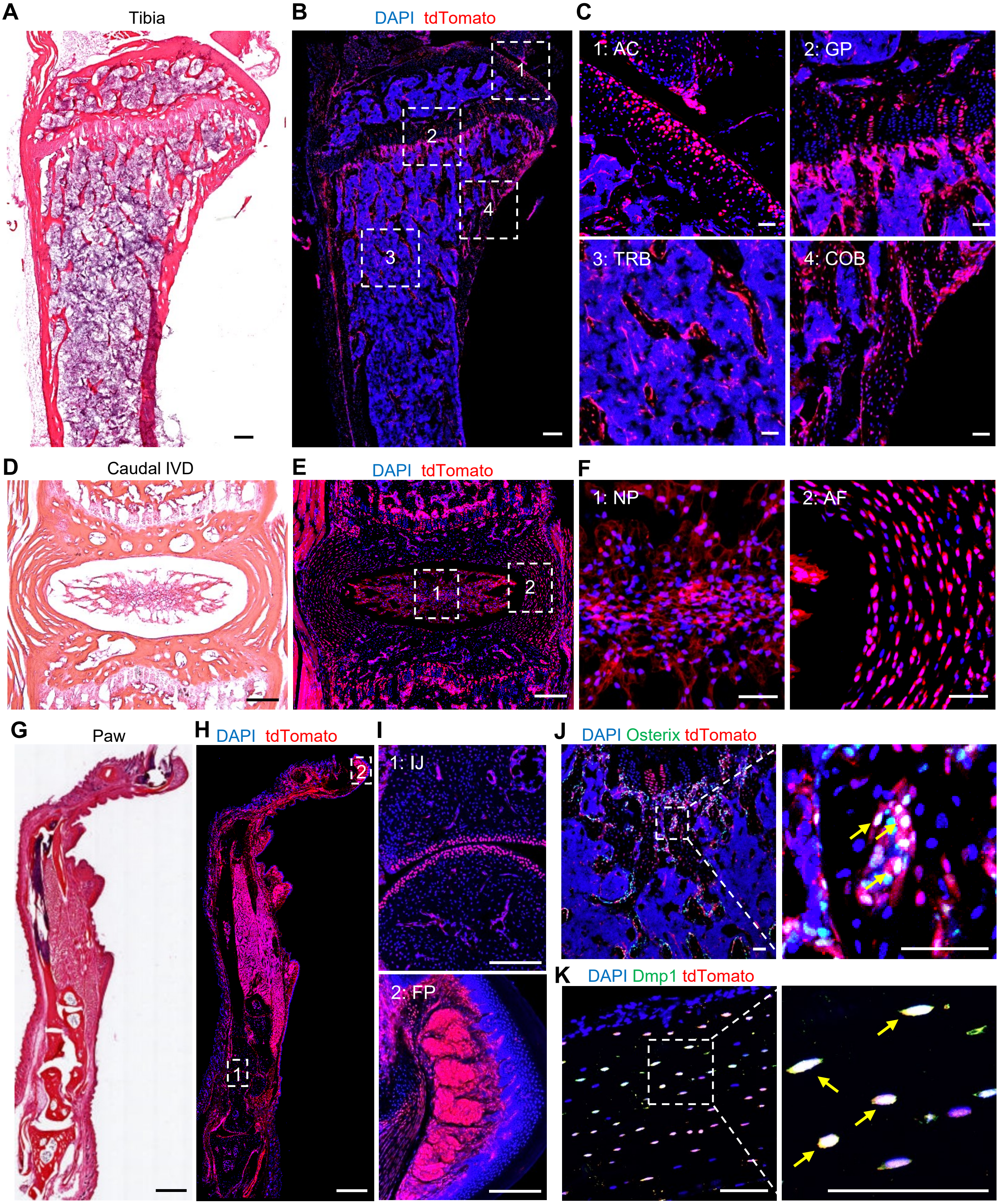

A
Tibia
B
DAPI tdTomato
1
2
3
4
C
1: AC
2: GP
3: TRB
4: COB
D
Caudal IVD
E
DAPI tdTomato
1
2
F
1: NP
2: AF
G
Paw
H
DAPI tdTomato
1
2
I
1: IJ
2: FP
J
DAPI Osterix tdTomato
K
DAPI Dmp1 tdTomato

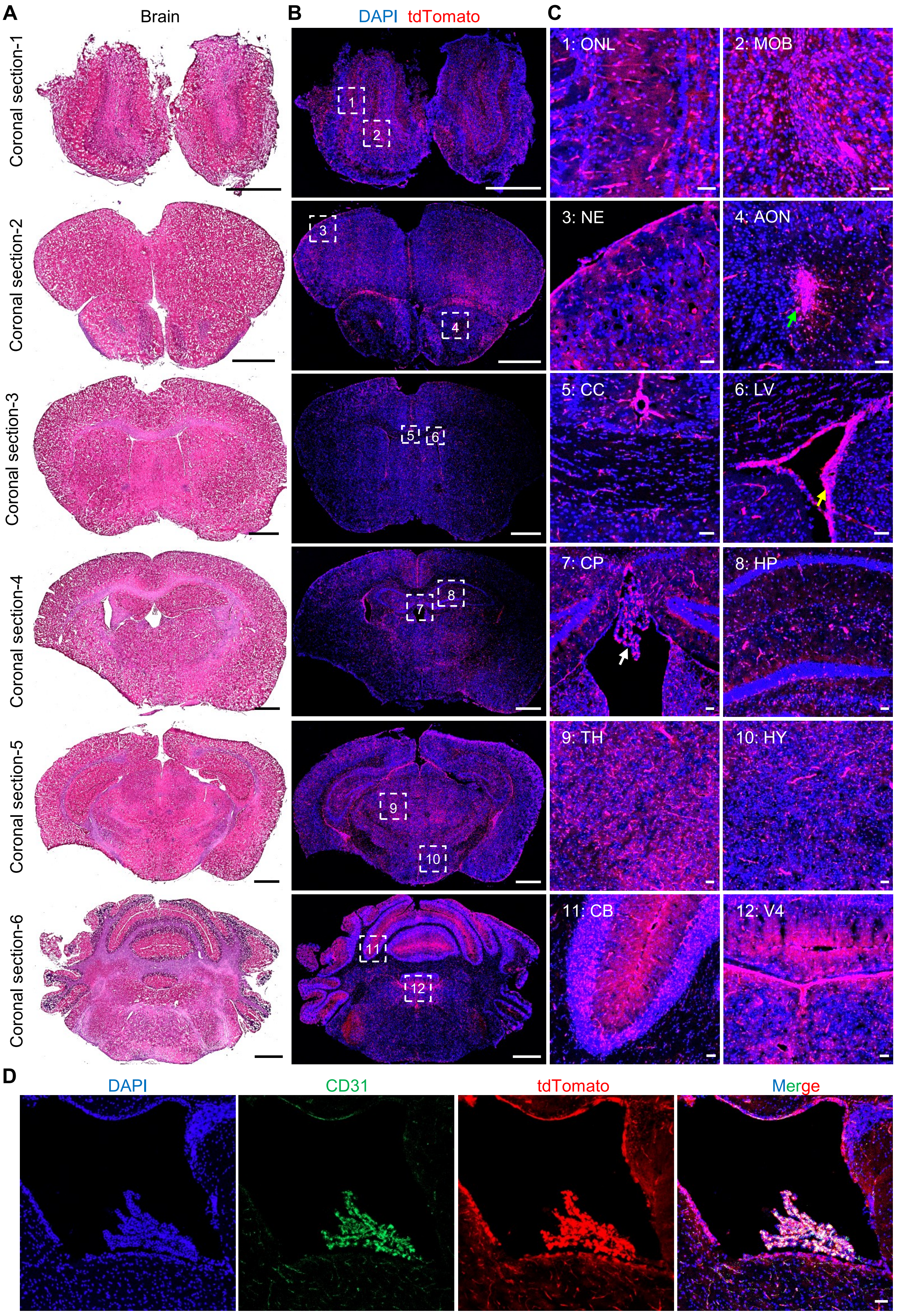
A
Brain
Coronal section-1
Coronal section-2
Coronal section-3
Coronal section-4
Coronal section-5
Coronal section-6
B
DAPI tdTomato
1
2
3
4
5
6
7
8
9
10
11
12
C
1: ONL
2: MOB
3: NE
4: AON
5: CC
6: LV
7: CP
8: HP
9: TH
10: HY
11: CB
12: V4
D
DAPI
CD31
tdTomato
Merge

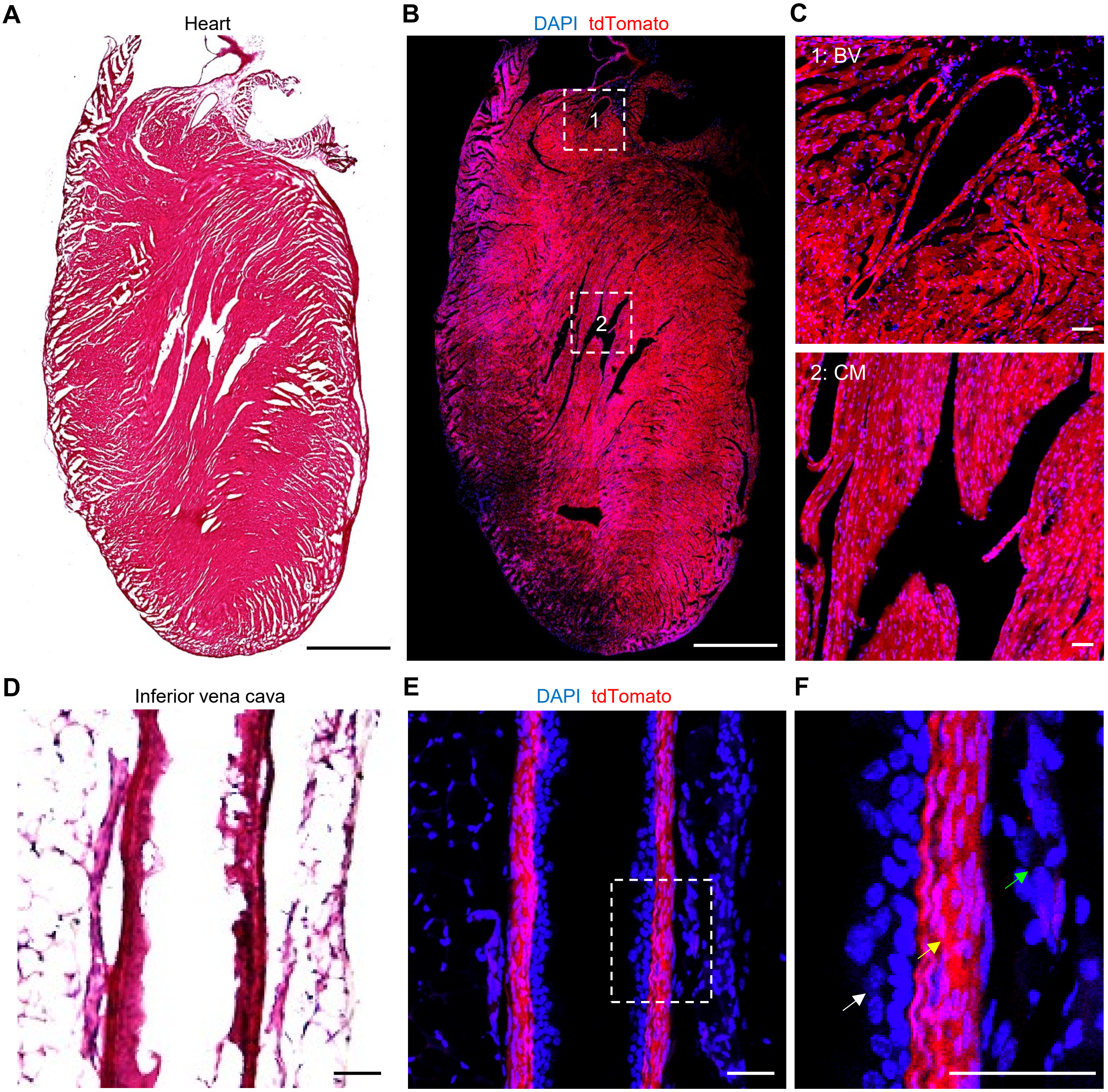
A
Heart
B
DAPI tdTomato
1
2
C
1: BV
2: CM
D
Inferior vena cava
E
DAPI tdTomato
F

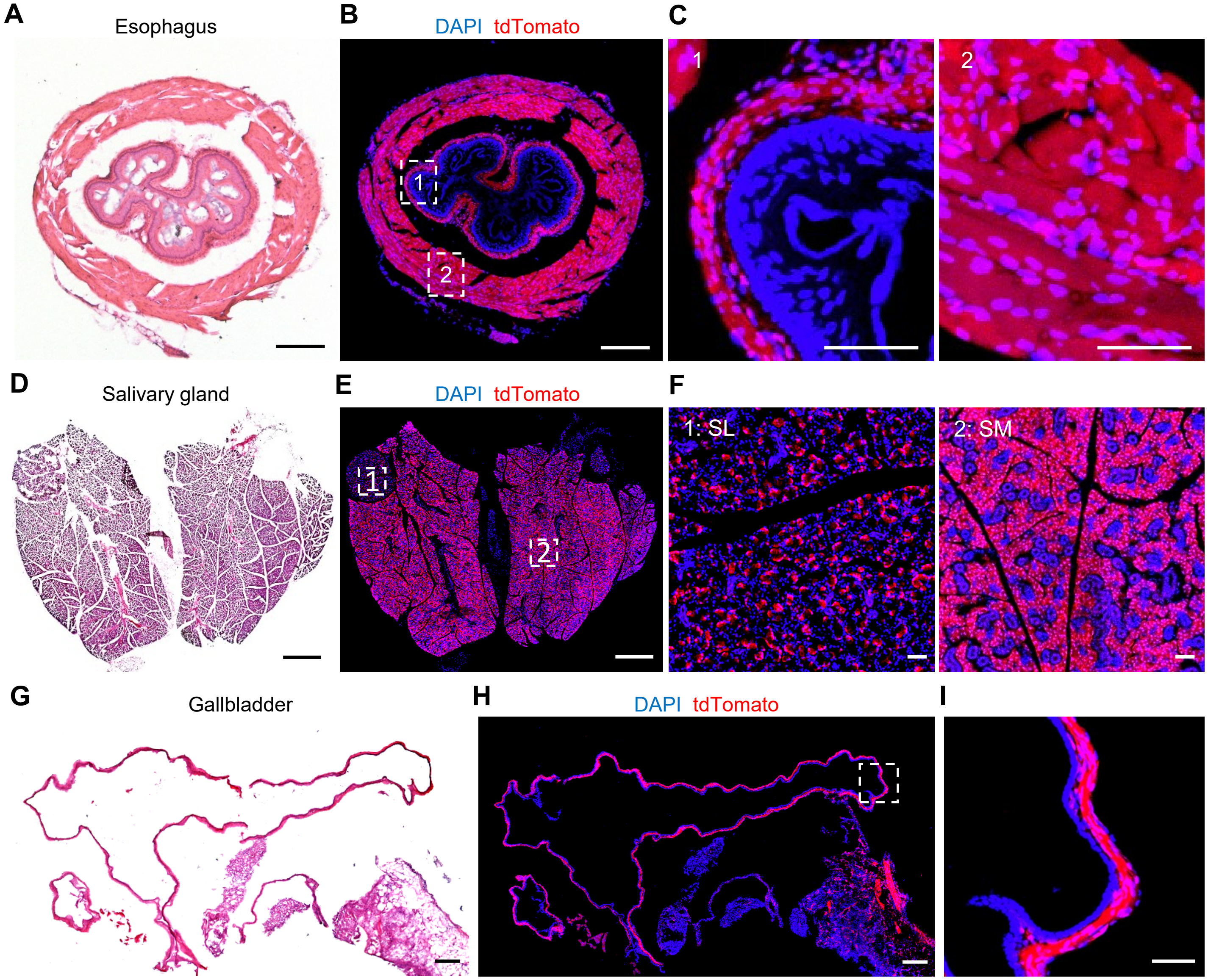

A
Esophagus
B
DAPI tdTomato
1
2
C
1
2
D
Salivary gland
E
DAPI tdTomato
1
2
F
1: SL
2: SM
G
Gallbladder
H
DAPI tdTomato
I

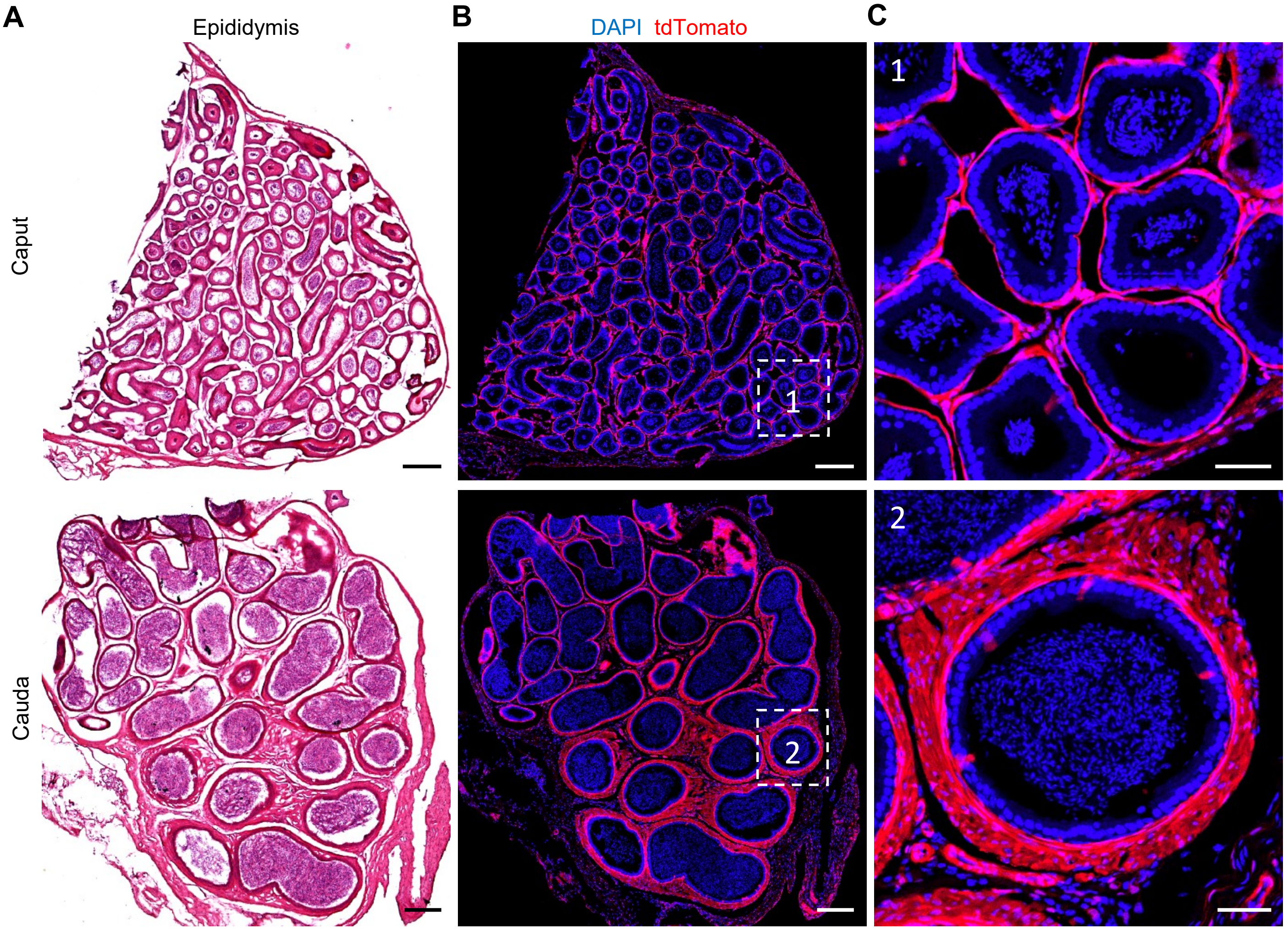
A
Epididymis
B
DAPI tdTomato
C
Caput
Cauda
1
2
1
2

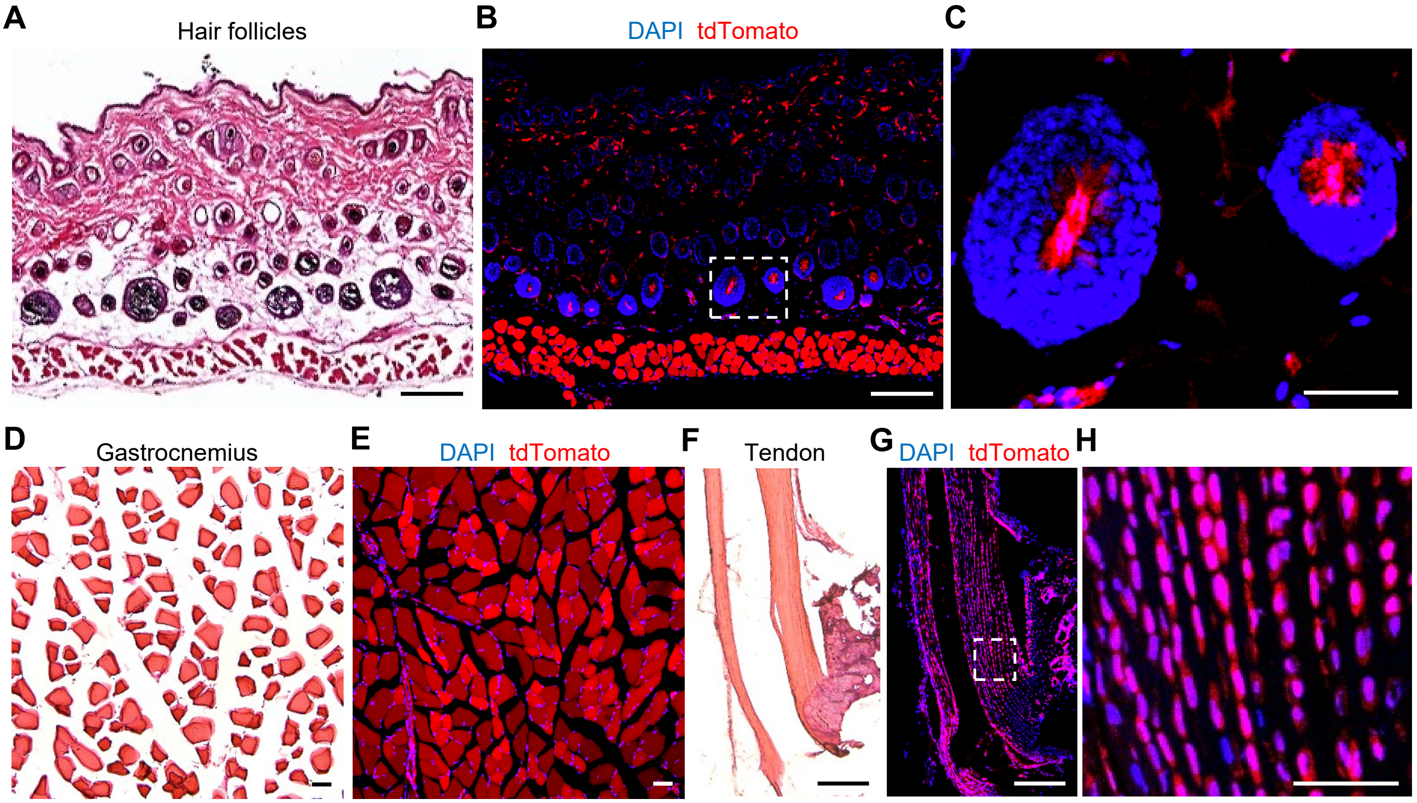
A
Hair follicles
B
DAPI tdTomato
C
D
Gastrocnemius
E
DAPI tdTomato
F
Tendon
G
DAPI tdTomato
H

Table S1. Comprehensive overview of Kindlin-2 expression across diverse tissues and organs

| Systems | Organs/Tissues | Subregion | Kindlin-2's expression |
|---|---|---|---|
| Skeletal system | Femur/Tibia | growth plate | ○ Partially positive |
| | | trabecular bone | ○ Predominantly localized to the trabecular bone surface |
| | | cortical bone | ○ Predominantly localized to osteocytes |
| | Knee joint | articular cartilage | ○ Primarily localized in the superficial zone of articular cartilage |
| | | meniscus | ○ Localized in the superficial zone of the meniscus |
| | | synovium | ○ Located on the synovial surface |
| | | subchondral bone | ○ Partially positive |
| | Intervertebral disc | cartilaginous endplate | ○ Partially positive |
| | | nucleus pulposus | ○ Overall positive |
| | | annulus fibrosus | ○ Partially positive |
| Nervous system | Brain | olfactory bulb | ○ Predominantly localized to cerebrovascular structures<br>○ Highly enriched in the choroid plexus |
| | | corpus callosum | |
| | | hippocampus | |
| | | neopallium | |
| | | thalamus | |
| | | hypothalamus | |
| | | cerebellum | |
| | | fourth ventricle | |
| | Spinal cord | gray matter | ○ Partially positive |
| | | white matter | ○ Partially positive |
| | Dorsal root ganglia | | ○ Partially positive |
| Respiratory system | Lung | bronchioli terminals | ○ Predominantly localized to non-epithelial compartments |
| | | alveolus pulmonis | ○ Overall positive |
| | Trachea | | ○ Predominantly localized within the cartilage layer |
| Circulatory system | Heart | blood vessel | ○ Overall positive |
| | | cardiac muscle | ○ Overall positive |
| | Aorta | | ○ Predominantly localized to the smooth muscle |
| | Inferior vena cava | | |

| | | | |
|---|---|---|---|
| | | | layer of the vascular tunica media |
| Digestive system | Salivary glands | sublingual gland | ○ Partially positive |
| | | submandibular gland | ○ Selectively enriched in acinar cells with no detectable expression in ductal structures |
| | Pancreas | pancreas islet | ○ Partially positive |
| | | pancreas acini | ○ Partially positive |
| | Liver | hepatocyte | ○ Overall positive |
| | | gallbladder | ○ Predominantly localized to the muscularis propria of the gallbladder |
| | Gastrointestinal tract | esophagus | ○ Predominantly expressed in the muscularis propria of the gastrointestinal tract and the lamina propria of the intestinal mucosa<br>○ Absent from the mucosal epithelium of the small intestine |
| | | stomach | |
| | | duodenum | |
| | | jejunum | |
| | | ileum | |
| | | cecum | |
| | | colon | |
| Urinary system | Kidney | glomerulus | ○ Predominantly localized to glomeruli and the renal medulla |
| | | collecting duct | |
| | | renal papillae | |
| | Bladder | mucosa | ○ Predominantly expressed in the muscularis propria of the urinary bladder |
| | | muscularis | |
| | Penis | urethra | ○ Absent from the urothelium |
| | | corpus cavernosum | ○ Partially positive |
| Reproductive system | Ovary | oocyte | ○ Virtually undetectable |
| | | oviduct | ○ Predominantly expressed in the muscularis of the oviduct |
| | Testis | seminiferous tubule | ○ Exclusively localized to the seminiferous tubule wall with no detectable expression in spermatogenic cells |
| | | tunica albuginea | ○ Overall positive |
| Immune system | Thymus | thymic cortex | ○ Partially positive |

| | | | |
|---|---|---|---|
| | | thymic medulla | ○ Partially positive |
| | | highly endothelial venule | ○ Overall positive |
| | | thymic capsule | ○ Virtually undetectable |
| | Spleen | white pulp | ○ Partially positive |
| | | red pulp | ○ Partially positive |
| | | central artery | ○ Overall positive |
| | | splenic capsule | ○ Overall positive |
| | Lymph node | superficial cortex | ○ Partially positive |
| | | paracortex | ○ Partially positive |
| | | medulla | ○ Partially positive |
| | | capsule | ○ Virtually undetectable |
| Other system | Eyeball | choroid | ○ Overall positive |
| | | retina | ○ Partially positive |
| | | crystalline lens | ○ Predominantly expressed in the lens epithelium |
| | | ciliary body | ○ Overall positive |
| | | cornea | ○ Partially positive |
| | Skin | surface epithelium | ○ Virtually undetectable |
| | | arrector pili muscle | ○ Overall positive |
| | | panniculus carnosus | ○ Overall positive |
| | Adipose tissue | brown adipose tissue | ○ Overall positive |
| | | white adipose tissue | ○ Overall positive |
